# Apparent Reconsolidation Interference Without Generalized Amnesia

**DOI:** 10.1101/2020.04.23.056952

**Authors:** Joaquín M. Alfei, Hérnan De Gruy, Dimitri De Bundel, Laura Luyten, Tom Beckers

**Affiliations:** Faculty of Psychology and Educational Sciences, KU Leuven, 3000 Leuven, Belgium; Leuven Brain Institute, KU Leuven, Leuven 3000, Belgium; Department of Biology, University of Rome, 185 Rome, Italy; Department of Pharmaceutical Sciences, Research Group Experimental Pharmacology, Center for Neurosciences, Vrije Universiteit Brussel, 1050 Brussels, Belgium

**Author notes:** Corresponding Author: Tom Beckers, Faculty of Psychology and Educational Sciences and Leuven Brain Institute, KU Leuven, Tiensestraat 102 box 3712, 3000 Leuven, Belgium. Both authors contributed equally to this work (Laura Luyten, Tom Beckers). E-mail addresses (J.M. Alfei), (H. De Gruy), (D. De Bundel), (L. Luyten), (T. Beckers).

**Keywords:** post-reactivation amnesia, memory reconsolidation, memory generalization, midazolam, ifenprodil

## Abstract

Memories remain dynamic after consolidation, and when reactivated, they can be rendered vulnerable to various pharmacological agents that disrupt the later expression of memory (i.e., amnesia). Such drug-induced post-reactivation amnesia has traditionally been studied in AAA experimental designs, where a memory is initially created for a stimulus A (be it a singular cue or a context) and later reactivated and tested through exposure to the exact same stimulus. Using a contextual fear conditioning procedure in rats and midazolam as amnestic agent, we recently demonstrated that drug-induced amnesia can also be obtained when memories are reactivated through exposure to a generalization stimulus (GS, context B) and later tested for that same generalization stimulus (ABB design). However, this amnestic intervention leaves fear expression intact when at test animals are instead presented with the original training stimulus (ABA design) or a novel generalization stimulus (ABC design). The underlying mechanisms of post-reactivation memory malleability and of MDZ-induced amnesia for a generalization context remain largely unknown. Here, we evaluated whether, like typical CS-mediated (or AAA) post-reactivation amnesia, GS-mediated (ABB) post-reactivation amnesia displays key features of a destabilization-based phenomenon. We first show that ABB post-reactivation amnesia is critically dependent on prediction error at the time of memory reactivation and provide evidence for its temporally graded nature. In line with the known role of GluN2B-NMDA receptor activation in memory destabilization, we further demonstrate that pre-reactivation administration of ifenprodil, a selective antagonist of GluN2B-NMDA receptors, prevents MDZ-induced ABB amnesia. In sum, our data reveal that ABB MDZ-induced post-reactivation amnesia exhibits the hallmark features of a destabilization-dependent phenomenon. Implication of our findings for a reconsolidation-based account of post-reactivation amnesia are discussed.

## 1. Introduction

Associative threat memories (also called fear memories) are at the heart of many anxiety-related disorders (Maddox, Hartmann, Ross, & Ressler, 2019). Historically, such memories have proven difficult to disrupt; even after seemingly successful interventions to attenuate the expression of fear memories, fearful reactions often reappear eventually, both in the lab (Urcelay & Miller, 2016) and in clinical treatment for anxiety (Vervliet, Craske, & Hermans, 2013). Over the last two decades, however, animal research focused on the neurobiology of Pavlovian fear conditioning has suggested a new avenue to dampen the expression of fear memories more permanently (Nader, Schafe, & Ledoux, 2000a, 2000b). Nader and collaborators (2000a) showed that re-exposure to training cues can reactivate a well-consolidated fear memory and render it vulnerable to disruption by inhibiting protein synthesis in the amygdala. As a result, the idea has gained ground that previously consolidated fear memories can, under certain conditions (e.g., Pedreira, Pérez-Cuesta, & Maldonado, 2004; Sevenster, Beckers, & Kindt, 2013), become deconsolidated or destabilized upon reactivation, in which state the memory will be vulnerable to pharmacological or other interventions that impair its retention and thus its later expression (resulting in amnesia). In other words, according to the canonical view, destabilized memories need to go through a protein synthesis-dependent process of reconsolidation for them to return to a stable form and be preserved (Lee, 2013).

In recent years, much has been learned regarding the mechanisms of reactivation-dependent destabilization (Zhang, Haubrich, Bernabo, Finnie, & Nader, 2018). In parallel, excitement has built over the prospect of reconsolidation-based interventions for psychological disorders in which emotional memories play a key role, including anxiety disorders, PTSD, mood disorders and substance abuse (for elaborate reviews, see Beckers & Kindt, 2017; Lee, Nader, & Schiller, 2017; Phelps & Hofmann, 2019). However, the vast majority of research on drug-induced amnesia (be it in humans or non-human animals) has made use of AAA experimental designs, where a memory is initially established for a cue or context A (CS) and later memory retrieval is produced or tested by exposure to the same CS. As such, research has largely ignored the conditions that may govern drug-induced post-retrieval amnesia beyond the original training situation (Beckers & Kindt, 2017). Yet this is a matter of considerable clinical relevance, for instance when trying to apply the principle of post-reactivation amnesia in clinical and sub-clinical samples where the learning history of maladaptive fear memories is usually unknown and where memories are rarely reactivated through exposure to the actual situation of learning (e.g., a combat war zone where a best friend was killed). Instead, clinical interventions typically rely on the use of generalization stimuli (GS) that bear a sufficient degree of similarity to the original event to reactivate the memory (e.g., war movie battle scenes) (for reviews, see Dymond, Dunsmoor, Vervliet, Roche, & Hermans, 2015; Walsh, Das, Saladin, & Kamboj, 2018).

We recently showed that the expression of a contextual fear memory, reactivated through exposure to generalization context B which is sufficiently similar to the original conditioning context A to elicit full fear generalization, could be disrupted through post-reactivation systemic administration of midazolam (MDZ), a positive allosteric modulator of the GABA_A_ receptor (Alfei et al., 2020). However, such amnesia was only evident upon later exposure to the same generalization context B (ABB, for training, reactivation and test context, respectively). No indication of MDZ-induced amnesia was observed when rats were instead exposed to the original learning situation (ABA) or to a different generalization context C (ABC) after reactivation. Like for other instances of post-retrieval amnesia (Alfei, Ferrer Monti, Molina, Bueno, & Urcelay, 2015; Ferrer Monti et al., 2017), sensitivity to amnestic effects of MDZ upon GS-mediated reactivation was driven by a temporal prediction error (PE) during reactivation: MDZ-induced amnesia after GS-mediated reactivation occurred only when there was a mismatch or discrepancy in US presentation between initial memory acquisition for context A (i.e., presentation of the US after 1 minute in A) and memory reactivation in context B (i.e., absence of the US or change in US timing during exposure to B).

The above-mentioned results may have important implications for the nature of post-retrieval amnesia. The fact that MDZ-induced amnesia is observed in an ABB procedure but not expressed in ABA or ABC procedures is difficult to reconcile with the canonical reconsolidation account of post-retrieval amnesia. According to that account, post-reactivation administration of an amnestic agent after memory destabilization (triggered by the occurrence of PE during exposure to context B) acts to prevent the protein-synthesis dependent return of the destabilized engram to a stable state, thus permanently and irreversibly disrupting the original fear memory trace (restorage deficit; Haubrich, Bernabo, Baker, & Nader, 2020). Such undoing of the engram should arguably impair memory expression regardless of whether animals are subsequently exposed to the generalization context B, the initial training context A, or a novel context C (Duvarci & Nader, 2004).

However, for the preserved expression of memory in ABA and ABC procedures to present a challenge to reconsolidation theory, it first needs to be demonstrated that ABB MDZ-induced amnesia indeed acts through similar mechanisms as AAA MDZ-induced amnesia (i.e., depends on the apparent destabilization of the contextual fear memory acquired during initial training in A). Alternatively, it has been proposed that impaired consolidation rather than impaired reconsolidation may account for ABB MDZ-induced post-retrieval amnesia. According to this idea, the presence of novel contextual elements when reactivating memory through exposure to B, while not detrimental to full retrieval of the fear memory acquired to A, might promote a switch from destabilization of the previously consolidated fear memory for A to the formation of a distinct fear memory representation (i.e., consolidation) for B that remains separate from the original memory representation (e.g., Hupbach, Hardt, Gomez, & Nadel, 2008). Building on this notion, it could be argued that an MDZ-based impairment in the consolidation of a fear memory for B rather than in the reconsolidation of the fear memory for A, is what produces a deficit in memory performance upon later test of B.

Before our earlier results can be accepted as a challenge for the reconsolidation account of post-retrieval amnesia, then, it remains to be demonstrated that the amnesia observed in an ABB procedure does indeed depend on apparent memory destabilization and does not reflect an impairment in consolidation of a new memory trace for the generalization context B. Here, to test these ideas, we use approaches that have been proposed to assess the canonical features of a destabilization-and-reconsolidation-based phenomenon (Nader & Hardt, 2009; Nader, Schafe, & Le Doux, 2000). These features are derived from the definition of memory destabilization as a transient memory state triggered by prediction error during memory reactivation, through which the original memory trace becomes malleable to be disrupted by amnestic interventions, and which requires a time-dependent process of restabilization (i.e., reconsolidation) for memory to persist (Elsey, Ast, & Kindt, 2018; Elsey & Kindt, 2017).

In the present series of experiments, we employed a contextual fear conditioning (CFC) procedure in rats to associate a context A with shock and then used a perceptually similar but discriminable context B for reactivation and test in order to evaluate fear memory generalization and amnesia. As a post-retrieval amnestic agent, we used MDZ, on the basis of previous reports demonstrating its ability to block contextual fear memory reconsolidation in rats (e.g., Bustos, Maldonado, & Molina, 2006, 2009). In Experiment 1, we replicated previous findings of our laboratory (Alfei et al., 2020) demonstrating that when a CFC memory for context A is reactivated through exposure to a perceptually similar context (B), subsequent MDZ administration attenuates fear responding to that same context B on a later test (ABB), but not fear responding to the initially trained context A (ABA). In further experiments, we show that MDZ has no such effect when administered outside the reconsolidation window (i.e., 6 h after memory reactivation; Experiment 2) and we dissociate immediate and delayed effects of post-reactivation MDZ administration, showing that ABB amnesia is expressed on a long-term memory test (24 h after reactivation) but no shortly after reactivation-plus-administration (Experiment 3). In Experiment 4, we show that administration of ifenprodil, a GluN2B-NMDA antagonist that has been demonstrated to block fear memory destabilization (Mamou, Gamache, & Nader, 2006), prior to memory reactivation, prevents MDZ-induced ABB post-retrieval amnesia without impairing expression of fear during reactivation. Finally, in Experiment 5, we show that MDZ does not disrupt the initial consolidation of a contextual fear memory. Collectively, our data suggest that MDZ-induced ABB amnesia displays the same cardinal characteristics of a destabilization-mediated phenomenon as other instances of post-retrieval amnesia.

## 2. Materials and Methods

See the figure legends for a detailed description of the designs and procedures of the experiments reported here.

### 2.1 Preregistration and data availability

Raw data for all the experiments reported here are available on the Open Science Framework at https://osf.io/zyx3p/. All designs, sample sizes, exclusion criteria and statistical analyses were preregistered on aspredicted.org. Preregistrations can be accessed through the following links: Experiment 1: http://aspredicted.org/blind.php?x=6ux9f9

Experiment 2: http://aspredicted.org/blind.php?x=ap7s8h

Experiment 3: http://aspredicted.org/blind.php?x=9xa63n3

Experiment 4: http://aspredicted.org/blind.php?x=4mw79m

Experiment 5: http://aspredicted.org/blind.php?x=3hn2r8

### 2.2 Subjects

Experimentally naive male Wistar rats (60-65 day old, weighing 290 - 340 g at the start of training) were obtained from Centro de Medicina Comparada (Esperanza, Santa Fe, Argentina). They were housed in groups of 4 in standard laboratory Plexiglas cages (60 cm long × 40 cm wide × 20 cm high) in a climate-controlled animal room in the Laboratorio de Psicología Experimental, Facultad de Psicología, Universidad Nacional de Córdoba (UNC), Argentina. For each experiment, new animals were obtained from the supplier and housed in the vivarium for at least 11 days before the start of the experiment to allow for acclimation. Food and water were available ad libitum. Animals were maintained on a 12-h light/dark cycle (lights on at 8 a.m.) at a room temperature of 21°. All procedures were approved by the animal ethics committees at UNC and KU Leuven and were in accordance with the Belgian Royal Decree of 29/05/2013 and European Directive 2010/63/EU.

### 2.3 Drugs and Administration

Midazolam (MDZ, Gobbi Novag SA, Buenos Aires, Argentina) was diluted in sterile isotonic saline (SAL, 0.9% w/v) to a concentration of 3 mg/ml and administered intraperitoneally (i.p.). The total volume of drug solution or equivalent amount of SAL was 1.0 ml/kg in all cases. Previous results have shown that this dose is effective for attenuating the retention of reactivated contextual fear memories in Wistar rats (Alfei et al., 2015; Ferrer Monti et al., 2017, 2016; Piñeyro, Ferrer Monti, Alfei, Bueno, & Urcelay, 2014). Ifenprodil (IFEN, Sigma-Aldrich Co.), a non-competitive, selective GluN2B-containing NMDA receptor antagonist, was dissolved in distilled water and administered intraperitoneally (i.p.) at a dose of 8.0 mg/kg, in a volume of 2.5 ml/kg for each rat (Rodrigues, Schafe, & Ledoux, 2001). Drug injections were always performed in a room that was different from the conditioning room.

### 2.4 Apparatus and Context Manipulation

Contextual Fear Conditioning (CFC) was conducted in a 250 × 250 × 250 mm chamber with 2 modular and removable grey opaque aluminum walls, a transparent Plexiglas ceiling and rear wall and a hinged front door (PanLab, Harvard Apparatus, US, controlled by PackWin V2.0 software). The floor consisted of 18 parallel stainless-steel rods, each measuring 4 mm in diameter, spaced 1.5 cm apart and connected to a device to provide adjustable footshocks (Unconditioned Stimulus, US). Variations in background odor, white noise, ventilation fan operation, house lights and wall color were employed to create two distinct physical contexts, A and B. Recording of behavior for offline analysis was done with digital video cameras mounted in front of the conditioning chamber. The chamber was enclosed in a sound-attenuating cubicle in a well-lit sound-attenuated experimental room.

**Context A.** For context A, the chamber contained a standard grid floor, uniformly colored walls and a white house light (three 5500 K – 6500 K LEDs with a 120° beam angle) mounted to the upper middle part of the left wall. A ventilation fan (65 dB) located in the upper back part of the cubicle was turned on and the chamber was cleaned with 80% ethanol prior to the session. The testing room light remained on throughout.

**Context B.** The alternative context contained the same standard grid floor, a grey opaque left wall and vertically striped (black/white) right wall. An infrared light (three bright 850 nm/940 nm wavelength LEDs with a 120° beam angle) mounted to the upper middle part of the right wall was turned on. The white house light and the ventilation fan were turned off. The chamber was cleaned with a household cleaning product prior to the session and the drop pan below the grid floor was scented with a thin layer of the same product. The testing room light remained on throughout.

### 2.5 Behavioral Procedures

In all experiments, rats were first individually labeled, weighed (one day before the start of handling and again on the fourth day of handling) and handled for 5 min on four separate days to habituate them to the experimenter. Handling was performed in a different room than the ones used for conditioning and drug administration. At the end of the second handling session, rats were injected with 1 ml/kg SAL, to habituate them to the injection procedure. Two contexts were created and used interchangeably as A (original context) and B (generalization context). The assignment of the different physical contexts to the roles of A and B was fully counterbalanced in all experiments. Transportation from the animal room to the experimental rooms always took place in a yellow plastic box filled with bedding and covered with a white cloth. All procedures were performed during the light phase of the diurnal cycle, between 10.00 am and 6.30 pm.

#### 2.5.1 Contextual fear conditioning (CFC)

Twenty-four h after the last day of handling, rats were taken individually from their home cage and transported to the conditioning chamber. The animals were exposed to context A for 1 min after which two footshocks (1.0 mA, 3-s duration, with an inter-shock interval of 30 s) were delivered. The total length of the CFC session was 1 min 36 s (Alfei et al., 2015; Ferrer Monti et al., 2017).

#### 2.5.2 Retrieval session (Experiments 1-4)

Memory retrieval always occurred 24 h after conditioning in context A. Rats were exposed to context B during 2 min without footshock for memory reactivation. MDZ 3 mg/kg or an equivalent amount of SAL was injected i.p. immediately after the reactivation. In Experiment 2, SAL or MDZ was additionally injected 6 h after reactivation. In Experiment 4, IFEN 8.0 mg/kg or an equivalent amount of SAL was injected i.p. 15 min prior to the reactivation session.

#### 2.5.3 Retention test

A memory retention test was carried out either 3.5 h (STM) or 24 h (LTM) after memory retrieval (except in Experiment 5). It consisted of a 5-min exposure to the chamber, without shock. Animals were either exposed to the same context as used for reactivation (ABB) or to the initial acquisition context (ABA). In Experiment 5, the retention test took place 24 h after memory acquisition, by exposing the animals to the same context as used for acquisition (AA).

### 2.6 Scoring of Freezing Behavior

In all experiments, freezing was used to index fear memory expression. It was defined as the total absence of body and head movements except for those associated with breathing. Given that previous reports have shown that software-scored freezing cannot reliably be compared across different contexts (Luyten, Schroyens, Hermans, & Beckers, 2014), freezing was scored manually, min-by-min, with a stopwatch and expressed as percentage of time. For all experiments, percentage of freezing per min during memory reactivation and the non-reinforced memory test was scored by two experienced raters blind to the experimental condition of each animal and inter-observer reliability was calculated (Pearson’s *r =* .95). Researchers were blinded to pharmacological treatments (but not physical contexts) during all behavioral testing procedures and fully blinded to group allocation during scoring of freezing behavior (all data files were randomized prior to scoring of freezing behavior).

### 2.7 Exclusion Criteria

For Experiments 1 - 4, the preregistered exclusion criterion stated that animals showing less than 20% of freezing during the reactivation session were considered as non-learners. For Experiment 5, this criterion was not set given that levels of freezing < 20 % during the retention test could have been caused by the post-acquisition injection of MDZ. Importantly, in all the experiments, excluded animals were replaced in order to obtain the pre-registered group sizes. All animals were given a number upon arrival in the lab, and replacement animals were included consecutively, following this numbering, until the prespecified sample size was reached. The exclusions are listed by experiment and group in Table 1, which can be accessed through the following link: https://osf.io/zyx3p/.

### 2.8 Statistical Analysis

Results are expressed as mean +/− the standard error of the mean (SEM) percentage of time the animals spent freezing. Data were analyzed by means of independent-samples t-tests or ANOVAs and effect sizes calculated as Cohen’s *d* (for t-tests) or *η^2^_p_* (ANOVAs). Significant one-way ANOVAs were followed up with Tukey tests, factorial ANOVAs with one-sided t-tests (MDZ < SAL) comparing MDZ versus SAL-treated animals per group. In case of deviation from normality, non-parametric Mann-Whitney U-tests were used rather than t-tests, and rank biserial correlation (*r_pb_*) was used to estimate effect sizes. In Experiments 3 and 5, MDZ treatment was expected to have no effect on freezing in the STM and retention test, respectively. A Bayesian independent-samples t-test was computed in each case (Bayes Factor _10_, with a default Cauchy prior width of *r* = 0.707) to gauge relative support for the null hypothesis versus the alternative hypothesis (in case of deviation from normality, a non-parametric Mann-Whitney U-test was computed and followed up with a Bayesian Mann-Whitney U Test). The Bayes factor (*BF*_10_) quantifies evidence in favor of the alternative hypothesis (H_A_; i.e., MDZ < SAL) relative to the null hypothesis (H_0_) (Jeffreys, 1961). A *BF*_10_ above 3 can be regarded as indicating substantial support in favor of the hypothesis in the nominator (H_A_) relative to the hypothesis in the denominator (H_0_), while values below .33 provide substantial evidence in favor of H_0_. In all cases, *p* < .05 was the statistical threshold. All analyses were carried out using JASP 0.9.0.1 (JASP, 2018) and all graphs were made with GraphPad Prism 7 (GraphPad Software Inc, La Jolla, CA, USA).

Of note, our statistical approach involved the evaluation of planned contrasts or specific interactions at test, based on a priori predictions (as recommended by Kirk, 1995, and Ruxton & Beauchamp, 2008, to preserve power in complex designs), instead of a more conservative, non-theory driven approach of conducting omnibus ANOVAs across reactivation and test and across all groups simultaneously and following up significant effects with post-hoc tests only. Our approach was afforded by the fact that we had clear directional hypotheses that were pre-specified in the preregistrations. Still, it bears mentioning that the pattern of significance across experiments might have looked slightly different using such a more conservative approach.

## 3. Results

### 3.1 Midazolam-induced amnesia for contextual fear does not generalize beyond the reactivation context

Previous results in our laboratory demonstrated that after fear memory acquisition for context A, contextual fear memory reactivation through exposure to a generalization context B can induce sensitivity to pharmacologically-induced amnesia for the generalization context (ABB), but this amnesia is not expressed when animals are later exposed to the originally trained context (ABA) (Alfei et al., 2020). In the present experiment, we sought to replicate this finding. We hypothesized that exposure to the generalization context would effectively retrieve the contextual fear memory for context A and that the subsequent generation of a PE through the absence of the US during memory reactivation would trigger memory vulnerability to the amnestic effects of MDZ, thus allowing MDZ-induced amnesia to be expressed upon a later test of the reactivation context (ABB).

The top panel of Fig. 1 presents an overview of the design. The bottom panel of Fig. 1 depicts contextual fear memory expression (freezing) during the reactivation and retention test. A one-way ANOVA on the reactivation data revealed no difference in freezing between the groups (*F*_(3,68)_ = .30, *p* = .82, *η^2^_p_* = .013). A factorial ANOVA on the retention test data (test context x drug treatment as factors) revealed a main effect of test context (*F*_(1,68)_ = 4.68, *p* = .034, *η^2^_p_* = .06), no main effect of drug treatment (*F*_(1,68)_ = 1.01, *p* = .318, *η^2^_p_* = .01), and a marginally significant interaction (*F*_(1,68)_ = 3.98, *p* = .050, *η^2^_p_* = .05). Planned comparisons indicated that MDZ produced an impairment in memory expression in the ABB-MDZ group relative to the ABB-SAL group, *t*_(34)_ = 2.52, *p* = .008, *d* = .81, whereas no effect was observed for the ABA groups, *t*_(34)_ = .82, *p* = .79, *d* = .27, suggesting that a difference between the initial training experience (i.e., US onset at the second min) and the reactivation session (i.e., US omission) triggered a period of memory vulnerability to MDZ (Alfei et al., 2015; Ferrer Monti et al., 2017), which resulted in amnesia when animals were later exposed to the reactivation context but not when they were exposed to the initially trained context.

**Fig. 1.**
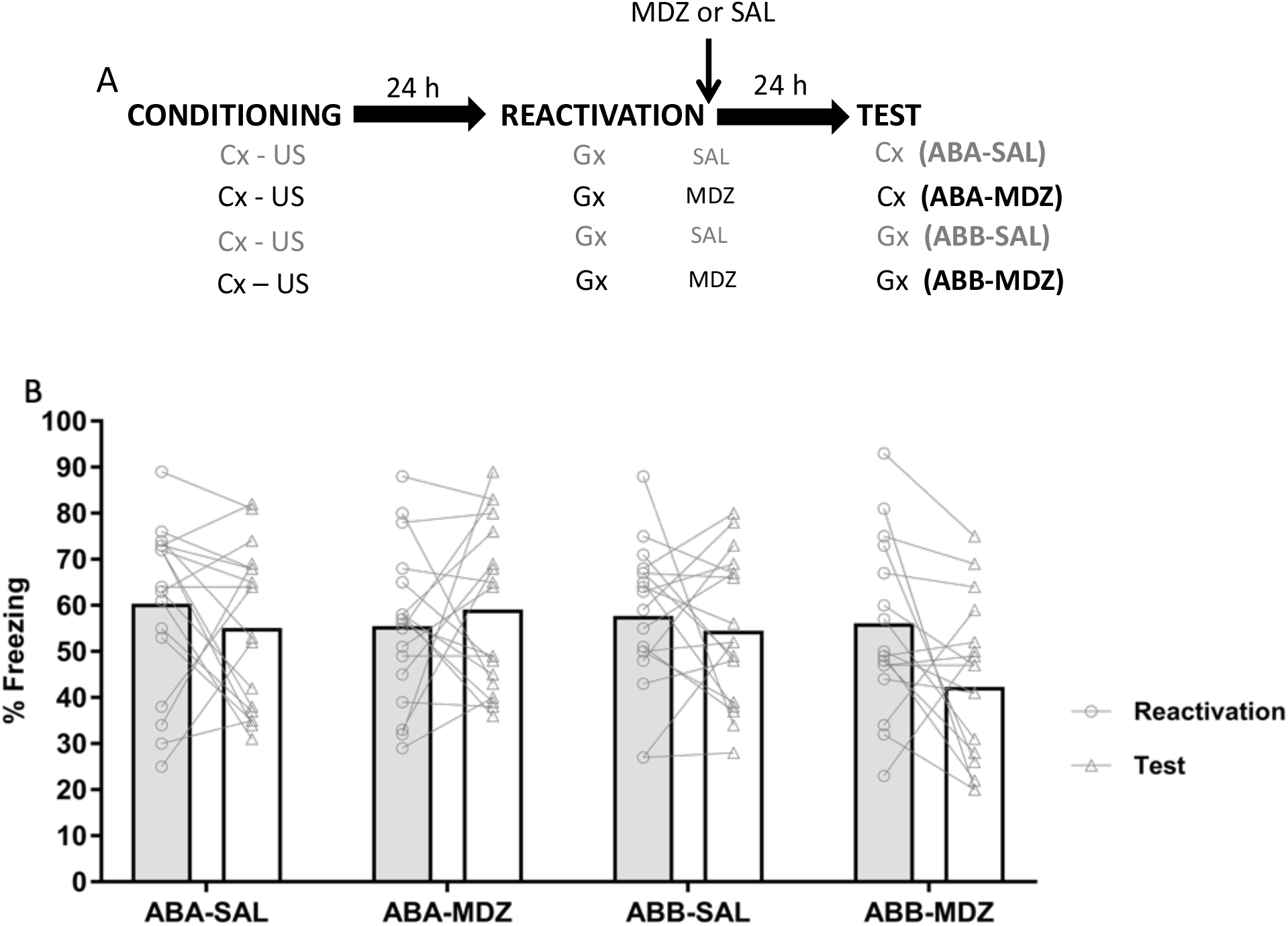
Midazolam-induced amnesia for contextual fear does not generalize beyond the reactivation context. **(A)** Four groups of rats received contextual fear conditioning for context A. Twenty-four h later, all groups received memory reactivation through exposure to context B, immediately followed by injection (i.p.) of 3 mg/kg MDZ or SAL (i.p.). Twenty-four h after memory reactivation, animals received a retention test in context A or context B. Which contexts served as A and B was fully counterbalanced. Animals were randomly assigned to groups (ABA-SAL: *n =* 18, ABA-MDZ: *n =* 18, ABB-SAL: *n =* 18, ABB-MDZ: *n =*18). **(B)** When memory retention was tested using the generalization context also used for memory reactivation, MDZ-induced amnesia was observed (i.e., a deficit in memory performance in the ABB-MDZ group relative to the ABB-SAL group). No differences in freezing were detected between the ABA-SAL and ABA-MDZ groups, indicating that experimentally induced amnesia was not obtained if animals were re-exposed to the initial training context at test. Data are expressed as means, symbols represent individual data points.

We thus replicate previous findings of our lab showing that a post-retrieval period of memory vulnerability to MDZ can indeed be triggered when a conditioned contextual fear memory is retrieved through exposure to a generalization situation. Critically, MDZ-induced attenuation of conditioned contextual fear responding is observed if the reactivation and test contexts are the same (ABB-MDZ) but fails to generalize to the originally trained context (ABA-MDZ) (Alfei et al., 2020). The subsequent experiments were designed to determine whether the post-retrieval memory impairment observed in the ABB-MDZ condition displays the defining features of a destabilization-and-reconsolidation-mediated phenomenon.

### 3.2 Midazolam-induced ABB amnesia is mediated by a time-limited process

According to memory consolidation theory, consolidation is a time-limited and protein-synthesis dependent process (Davis & Squire, 1984): protein synthesis processes that occur within a 4-6 h time window after encoding are critical for long-term retention. It is widely accepted that reconsolidation is likewise time-limited: When a memory trace becomes active and labile upon retrieval (memory reactivation), its restabilization is assumed to take a limited amount of time (Nader & Hardt, 2009). Traditionally, this notion is based on the observation that administration of MDZ and other amnestic agents will result in a long-lasting decrement in memory performance only if the amnestic agent is administered within a limited time window following memory reactivation (i.e., within a period of up to 6 h after memory destabilization; Nader et al., 2000a). The “reconsolidation window” refers to the fact that the amnestic gradient decreases with the interval between reactivation and the amnestic procedure (Przybyslawski & Sara, 1997). Importantly, if the post-retrieval attenuation that is observed when a memory acquired for context A is retrieved and tested using a generalization context B (ABB design) is mediated through a similar plasticity mechanism (Fig. 1, ABB-MDZ group), one should reasonably expect that increasing the interval (≥6 hours) between fear memory reactivation and MDZ injection should likewise abolish the ABB-MDZ amnestic effect. Experiment 2 was designed to test this hypothesis.

The top panel of Fig. 2 presents an overview of the design. The bottom panel of Fig. 2 depicts memory performance during reactivation and the retention test. An independent-samples t-test on reactivation data revealed no difference between the groups, *t*_(32)_ = .52, *p* = .60, *d* = .17. A one-sided t-test (MDZ-SAL < SAL-MDZ) showed that animals that were treated with MDZ immediately after reactivation and SAL 6 h later expressed significantly less freezing on the retention test than animals treated with SAL immediately after reactivation and MDZ 6 h later, *t*_(32)_ = 2.44, *p* = .01, *d* = .83.

**Fig. 2.**
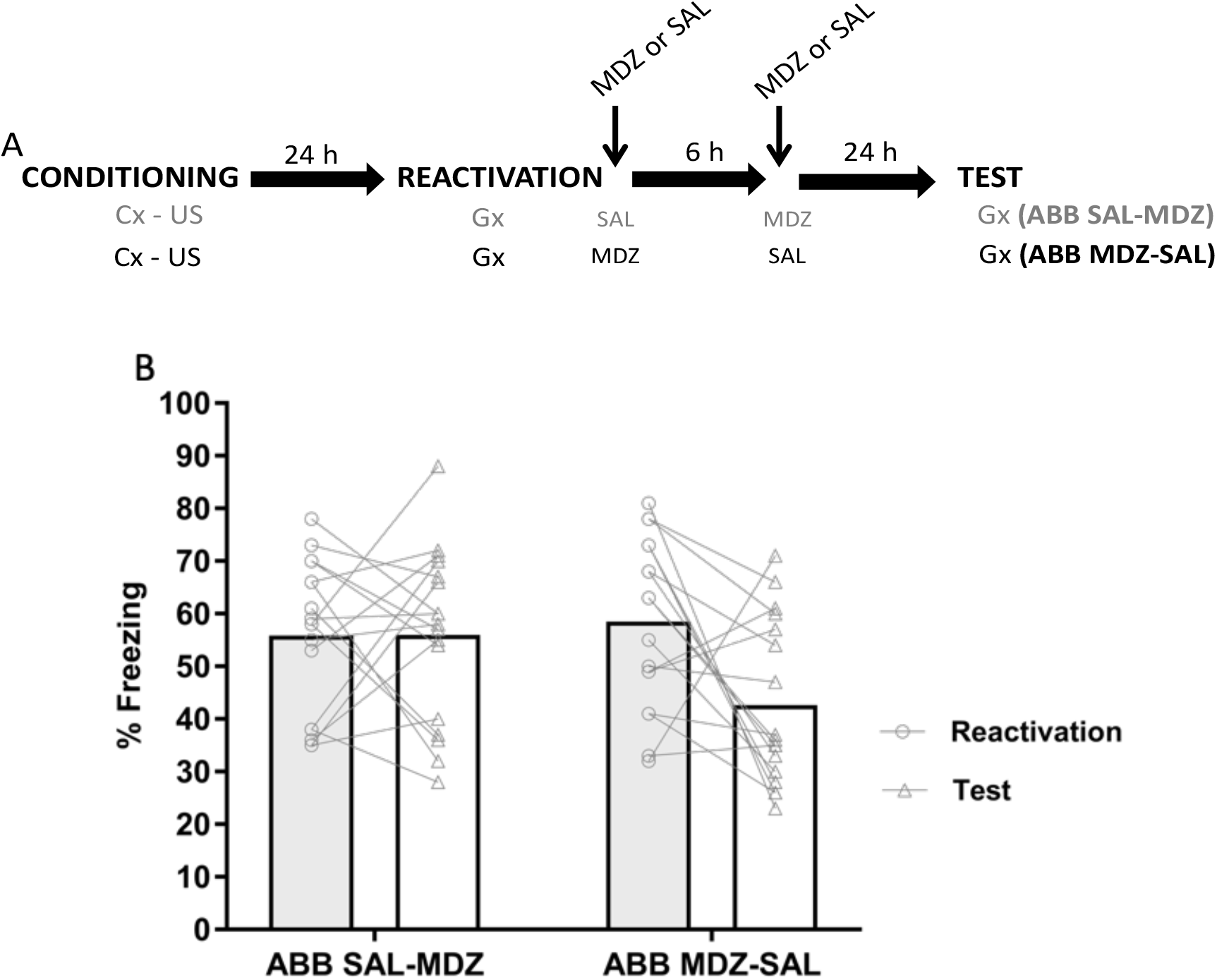
Midazolam-induced ABB amnesia is mediated by a time-limited process. **(A)** wo groups of rats received contextual fear conditioning for context A. Twenty-four h later, all groups received memory reactivation through exposure to context B. The first group received injections (i.p.) of SAL immediately after memory reactivation and of 3 mg/kg MDZ 6 h later. The second group was injected (i.p) with 3mg/kg of MDZ immediately after memory reactivation and SAL (i.p.) 6 h later. All groups were subjected to a retention test in context B 24 h after memory reactivation. Which contexts served as A and B was fully counterbalanced. Animals were randomly assigned to groups (SAL-MDZ: *n =* 18, MDZ-SAL: *n =* 18). **(B)** Rats that received MDZ immediately after memory reactivation and SAL 6 h later displayed a significant impairment in memory expression relative to rats that received SAL immediately after memory reactivation and MDZ 6 h later. Data are expressed as means, symbols represent individual data points.

Our data suggest that extending the time interval between contextual fear memory reactivation through exposure to a generalization context, and the injection of MDZ, prevents reactivation-mediated ABB amnesia that is observed when MDZ is administered immediately after memory reactivation (see also Fig. 1, ABB-MDZ group). Moreover, given that MDZ was administered in both groups, and even more closely to the retention test in the SAL-MDZ group than in the MDZ-SAL group, residual activity of the drug at the time of the retention test cannot account for the reduction in fear expression observed in the MDZ-SAL group. Overall, our results are consistent with the notion that memory destabilization confers a time-limited sensitivity to amnestic interventions and the hypothesis that MDZ-induced ABB amnesia is a destabilization-mediated phenomenon (Nader & Hardt, 2009).

### 3.3 Midazolam-induced ABB amnesia is expressed on a long-term memory test but not on a short-term memory test

The temporally graded nature of memory reconsolidation is also evident from the fact that when memory reactivation is followed by the administration of an amnestic drug, memory performance is typically intact on an immediate, short-term memory test (e.g., 6 h after the amnestic intervention) (Debiec, Doyere, Nader, & LeDoux, 2006; Duvarci & Nader, 2004; Ponnusamy et al., 2016). It is only on a long-term memory test (e.g., 24 h later) that reconsolidation interference may result in amnesia (Nader, Schafe, & Ledoux, 2000a). This suggests that it is indeed memory restabilization rather than memory expression that is affected by the amnestic treatment. Accordingly, if the post-retrieval memory decrement that we observe in our ABB-MDZ group is indeed a reconsolidation-mediated phenomenon, we should expect it to be absent on a short-term memory test (STM) and a performance deficit should be expressed on a long-term memory test (LTM) only. Testing this hypothesis was the aim of Experiment 3.

The top panel of Fig. 3 presents an overview of the design. The bottom panel shows the results. A one-way ANOVA on the reactivation data revealed no differences in freezing between the groups (*F*_(3,68)_ = .09, *p* = .96, *η^2^_p_* = .004). On the STM test, no differences in freezing were observed between the MDZ and SAL groups, *u* = 152, *p* = .76, *r_pb_* = .06. A Bayesian Mann-Whitney U Test yielded substantial evidence for the absence of a group difference, *BF*_10_ = .33. In contrast, on the LTM test, animals in the MDZ group exhibited less freezing than those in the SAL group, *t*_(34)_ = 2.25, *p* = .031, *d* = .75. The intact freezing in MDZ-treated animals on the STM test combined with impaired performance in MDZ-treated animals on a LTM test suggests that MDZ administration following exposure to generalization context B affected a reconsolidation-based process (Duvarci & Nader, 2004).

**Fig. 3.**
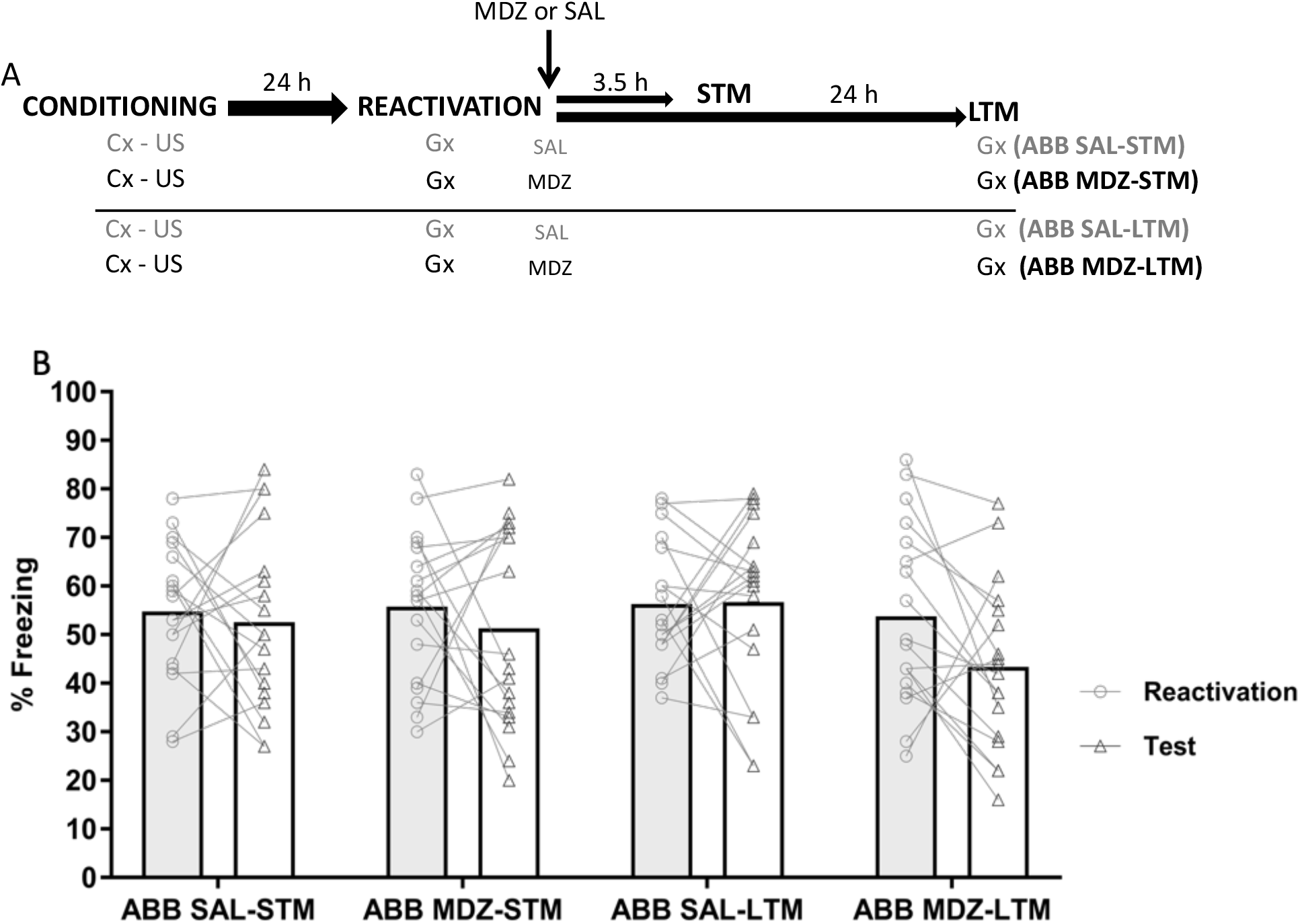
Midazolam-induced ABB amnesia is expressed on a long-term memory test but not on a short-term memory test. **(A)** Four groups of rats received CFC for context A. Twenty-four h later, all groups received memory reactivation through exposure to generalization context B. Immediately after reactivation, half of the animals were injected with MDZ (3 mg/kg, i.p.) whereas the other half were injected with SAL (i.p.). Three and a half h after the reactivation session, one of the MDZ and one of the SAL groups were given a short-term memory (STM) retention test involving context B. The other 2 groups received a long-term memory (LTM) retention test involving context B 24 h after memory reactivation. Which contexts served as A or B was fully counterbalanced. Animals were randomly assigned to groups (MDZ-STM: *n =* 18, SAL-STM: *n =* 18, MDZ-LTM: *n =* 18, SAL-LTM: *n =* 18). **(B)** Relative to SAL controls, rats injected with MDZ after memory reactivation using generalization context B showed an impairment on a LTM test involving the same context B, in line with the results of Exp 1 and 2. Rats given a STM test instead of a LTM test did not show a significant deficit in performance relative to controls. Data are expressed as means, symbols represent individual data points.

### 3.4 Ifenprodil administration prior to memory reactivation prevents MDZ-induced ABB amnesia

Memory consolidation and reconsolidation are both inferred from the temporally graded nature of amnesia (Dudai, 2004). As such, the features of the ABB MDZ-induced amnesia observed in Exp. 1-3 are not exclusively compatible with a destabilization-based phenomenon. As noted in the introduction, ABB MDZ-induced amnesia could also be accounted for by a consolidation-based mechanism. In the next two experiments we evaluate this possibility.

Several studies support the view that memory destabilization and its subsequent reconsolidation involve the activity of different subtypes of glutamate receptors (NMDAR) in the basolateral complex of the amygdala (BLA). Particularly, there is evidence that GluN2B-NMDARs are critically involved in fear memory destabilization, whereas GluN2A-NMDARs are recruited for restabilization (Milton et al., 2013; Wang, De Oliveira Alvares, & Nader, 2009). In line with this view, intra-BLA infusions of ifenprodil (IFEN), a GluN2B-NMDAR antagonist, leave memory reactivation and reconsolidation intact but prevent memory destabilization of cued (e.g., Mamou et al., 2006; Milton et al., 2013) and contextual fear memories (e.g., Ferrer Monti et al., 2016; Haubrich et al., 2015). Furthermore, IFEN administration prior to initial memory acquisition or extinction learning has no acute effect on the expression of fear but disrupts the consolidation process (Bauer, Schafe, & Ledoux, 2002; Blair, Sotres-bayon, Moita, & Ledoux, 2005; Rodrigues et al., 2001; Sotres-bayon, Bush, & Ledoux, 2007).

To corroborate the hypothesis that MDZ-induced ABB amnesia as observed in Experiment 1-3 is mediated by destabilization of the contextual fear memory established for A, rather than disrupted consolidation of a contextual fear memory for B formed during reactivation, we investigated the involvement of GluN2B-NMDAR in MDZ-induced ABB amnesia. To this end, we evaluated the effect of systemic injection (i.p.) of IFEN prior to memory reactivation on MDZ-induced ABB amnesia. If MDZ-induced ABB amnesia depends on a destabilization process, IFEN given before memory reactivation in B should block the post-retrieval amnestic effects of MDZ since the memory trace should not destabilize.

The top panel of Fig. 4 presents an overview of the design. The bottom panel of Fig. 4 depicts memory performance during reactivation and the retention test. A factorial ANOVA on the reactivation data (pre-treatment x post-treatment as factors) revealed no main effect of pre-treatment (*F*_(1,68)_ = .02, *p* = .88, *η^2^_p_* = .001), no main effect of post-treatment (*F*_(1,68)_ = .09, *p* = .75, *η^2^_p_* = .001), and no pre-treatment x post-treatment interaction (*F*_(1,68)_ = .04, *p* = .83, *η^2^_p_* = .001). So, in line with previous studies using systemic administration (Rodrigues et al., 2001; Sotres-Bayon, Bush, & Ledoux, 2007) or BLA infusion (Ferrer Monti et al., 2016; Milton et al., 2013), IFEN did not acutely affect the behavioral expression of contextual fear memory compared to SAL (Fig. 4B). A factorial ANOVA on the retention test data (pre-treatment x post-treatment as factors) revealed a main effect of pre-treatment (*F*_(1,68)_ = 4.76, *p* = .03, *η^2^_p_* = .06), no effect of post-treatment (*F*_(1,68)_ = 2.48, *p* = .12, *η^2^_p_* = .03), and a trend towards a pre-treatment x post-treatment interaction (*F*_(1,68)_ = 3.31, *p* = .07, *η^2^_p_* = .04). Planned comparisons revealed lower freezing in the MDZ than the SAL post-treatment group when SAL was administered prior to memory reactivation, *t*_(34)_ = 2.29, *p* = .01, *d* = .76, but not when animals received IFEN prior to memory reactivation, *t*_(34)_ = .18, *p* = .57, *d* = .06. Moreover, SAL-MDZ-treated animals showed significantly less freezing than IFEN-MDZ treated animals, *t*_(34)_ = 2.82, *p* = .004, *d* = .87 (Fig. 4B), suggesting that the effect of post-reactivation MDZ administration on later fear memory expression is prevented by blockade of GluN2B-NMDA-dependent memory destabilization. The results of this experiment suggest that systemic administration of the GluN2B-NMDAR antagonist IFEN prevents the destabilization of contextual fear memory acquired for A through exposure to generalization context B.

**Fig. 4.**
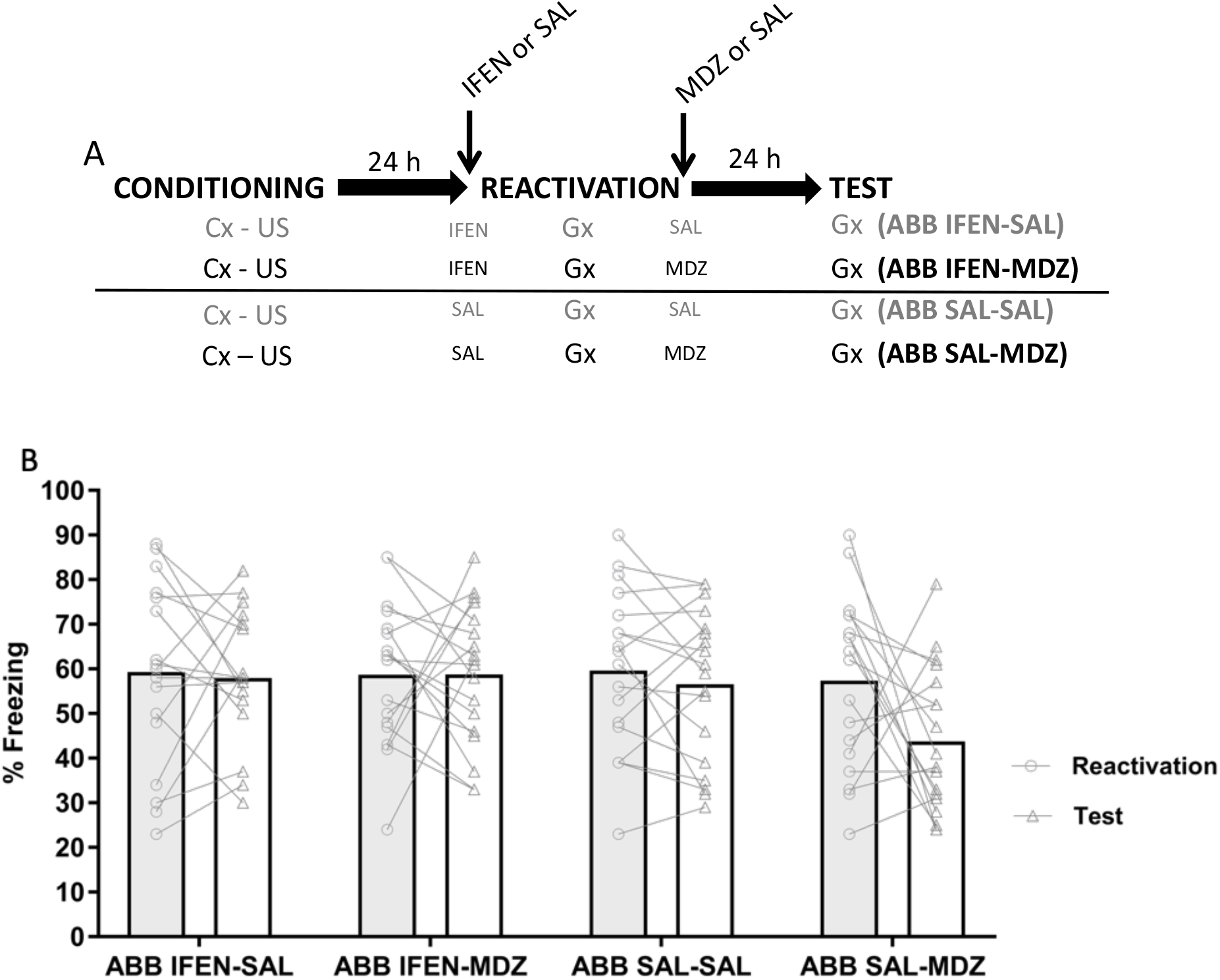
Ifenprodil administration prior to memory reactivation prevents MDZ-induced ABB amnesia. **(A)** Four groups of rats received CFC for context A. Twenty-four h later, half of the rats were injected with IFEN (8.0 mg/kg, i.p.) whereas the other half was injected with SAL (i.p.). Fifteen minutes later, animals were subjected to fear memory reactivation through exposure to context B and immediately afterwards injected with either SAL (i.p.) or MDZ (3 mg/kg, i.p.). All groups were given a memory retention test for context B 24 h after memory reactivation. Which contexts served as A or B was fully counterbalanced. Animals were randomly assigned to groups (IFEN-SAL: *n =* 18, IFEN-MDZ: *n =* 18, SAL-SAL: *n =* 18, SAL-MDZ: *n =* 18). **(B)** Systemic administration of the GluN2B-selective NMDAR antagonist IFEN has no effect on the expression of generalized contextual fear memory (i.e., similar levels of freezing across groups during memory reactivation). Conditioned freezing behaviour during test by group reveals that IFEN blocks MDZ-induced ABB amnesia. No differences were detected between IFEN-SAL and IFEN-MDZ, whereas SAL-MDZ animals exhibited reduced fear memory expression relative to SAL-SAL controls. Data are expressed as means, symbols represent individual data points.

As mentioned above, pre-training administration of IFEN has been shown to impair fear memory consolidation. Therefore, if MDZ-induced ABB amnesia were mediated by a consolidation process rather than a destabilization-based process, one should expect a memory impairment at test in animals that received IFEN prior to memory reactivation (i.e., less freezing in the IFEN-SAL group than the SAL-SAL group). However, a (not preregistered) independent-samples t-test did not yield a significant difference in memory expression between IFEN-SAL and SAL-SAL animals, *t*_(34)_ = .25, *p* = .80, *d* = .08. The lack of a memory deficit at test in the IFEN-SAL group again suggests that the MDZ effects reported above are likely due not to a consolidation-mediated but to a destabilization-mediated process.

### 3.5 Midazolam does not interfere with the consolidation of contextual fear memory

In a final experiment, to further corroborate the hypothesis that MDZ-induced post-retrieval amnesia reflects destabilization-based memory malleability rather than a disrupted consolidation process, we examined whether post-training administration of MDZ disrupts the consolidation of a newly encoded contextual fear memory. The result of this experiment should help to illuminate the process underlying the MDZ-induced ABB amnesia observed in the previous experiments. If systemic injection of MDZ applied shortly after the acquisition of a new contextual fear memory impairs the consolidation (and thus, the later expression) of that contextual fear memory, it would be reasonable to consider that ABB MDZ-induced post-retrieval amnesia originates from MDZ blocking the consolidation of a newly formed fear memory for the generalization context B.

The top panel of Fig. 5 presents an overview of the design. The bottom panel of Fig. 5 depicts memory performance during the retention test. The level of freezing of MDZ-treated animals was not significantly less than that of their SAL-treated counterparts on the retention test, *t*_(36)_ = .30, *p* = .61, *d* = .09. A Bayesian analogue one-sided t-test yielded substantial evidence for the absence of a group difference, *BF_10_* = .25, suggesting that a 3 mg/kg dose of MDZ does not significantly disrupt the consolidation of a contextual fear memory.

**Fig. 5.**
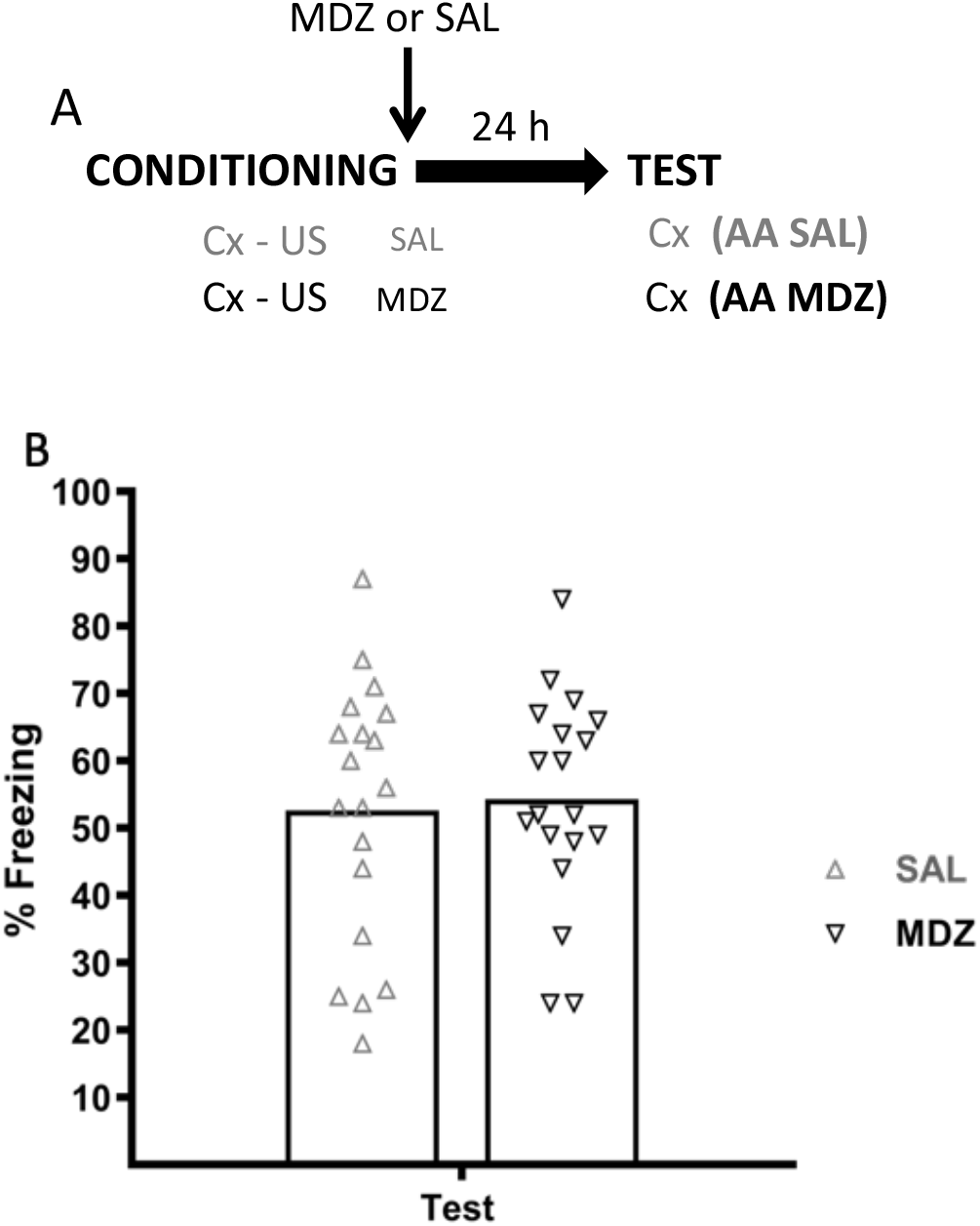
Midazolam does not interfere with the consolidation of contextual fear memory. **(A)** Two groups of rats received CFC for context A. Immediately after training, half of the animals were injected with MDZ (3 mg/kg, i.p.), whereas the other half received SAL (i.p.). Both groups were given a retention test in context A 24 h later. Animals were randomly assigned to groups (SAL: *n =* 19, MDZ: *n =* 19). **(B)** Conditioned contextual fear expression (i.e., freezing) at test was not significantly affected when MDZ was administered shortly after acquisition. Data are expressed as means, symbols represent individual data points.

In line with published studies using contextual fear memories and post-acquisition systemic injection of MDZ (3 mg/kg dose or lower) (Bustos et al., 2006; Pain, Launoy, Fouquet, & Oberling, 2002), our results reveal that MDZ does not disrupt contextual fear memory when it is administered shortly following the training session (i.e., MDZ-treated animals do not differ from SAL group). This suggests that disrupted consolidation cannot be held responsible for the observed amnestic effects of MDZ administration after fear memory reactivation though exposure to a generalization context, and adds further support to the idea that the MDZ-induced ABB amnesia observed in Experiments 1-4 reflects a process similar to that involved in other “reconsolidation interference” experiments.

## 4. Discussion

We previously demonstrated that post-retrieval amnesia for contextual fear memory can be observed when memory reactivation is achieved through exposure to a generalization context B prior to MDZ administration and fear is later tested for the same context B (ABB), but MDZ-induced post-retrieval amnesia is not obtained when animals are instead exposed to the original training context (ABA) or an equally similar but discriminable context C (ABC) at test (Alfei et al., 2020). The aim of the current paper was to resolve a number of open questions regarding the underlying mechanisms of such MDZ-induced ABB amnesia. Collectively, the present results suggest that MDZ-induced ABB post-retrieval amnesia reflects a destabilization-mediated form of memory malleability.

Replicating previous findings of our laboratory, we first showed that contextual fear memory can be rendered sensitive to interference through exposure to a generalization context (also see Kroes, Dunsmoor, Lin, Evans, & Phelps, 2017; Soeter & Kindt, 2015), but the resultant MDZ-induced amnesia can be observed only when reactivation and test contexts are the same (Fig. 1; ABB group). Given that consolidation and reconsolidation processes are both inferred from evidence for a transient period of memory vulnerability (Miller & Matzel, 2000), a straightforward prediction of both accounts of experimental amnesia is that an amnestic agent should lead to a long-lasting memory deficit only if administered soon after training or reactivation. Indeed, we demonstrated that post-retrieval ABB amnesia is observed if MDZ is given immediately after CFC reactivation, but not when MDZ administration is delayed for 6 h (Fig. 2), indicating that the interaction between reactivation and MDZ is dependent on the drug being administered within a specific time window (Nader, Schafe, & Ledoux, 2000a). Furthermore, post-reactivation administration of MDZ, while effective in disrupting LTM performance for the generalization context B, did not affect STM fear expression for context B (Fig. 3) (Duvarci & Nader, 2004). The critical requirement of memory destabilization for MDZ-induced ABB post-retrieval amnesia was demonstrated in Experiment 4: MDZ-induced ABB amnesia was prevented when IFEN was administered prior to memory reactivation (Fig. 4) (Milton et al., 2013), which contradicts the notion that MDZ-induced ABB amnesia reflects a deficit in consolidation. Experiment 5 added further support to the claim that MDZ-based ABB amnesia is a destabilization rather than a consolidation-based phenomenon by showing that MDZ at the dose used here is unable to disrupt the consolidation of a contextual fear memory (Fig. 5) (Bustos et al., 2006).

As noted in the introduction, the canonical reconsolidation theory of post-retrieval amnesia assumes that consolidated fear memories, which are in principle insensitive to amnestic interventions, can revert to a vulnerable state if they are retrieved under conditions that involve an appropriate degree of prediction error (for elaborate reviews of the literature, see Exton-McGuinness, Lee, & Reichelt, 2015; Fernández, Boccia, & Pedreira, 2016; Krawczyk, Fernández, Pedreira, & Boccia, 2017; Lee, 2013; Sinclair & Barense, 2019). The engagement of memory destabilization-and-reconsolidation is assumed to be mediated by protein synthesis-dependent plasticity of the original memory trace (Morris et al., 2006), thereby allowing post-retrieval manipulations that block protein synthesis to prevent restorage of the original trace (Khalaf et al., 2018). A straightforward prediction of the reconsolidation account of experimentally induced amnesia is that exposure to a generalization context that effectively and fully retrieves a previously acquired contextual fear memory should be able to render the original memory representation sensitive to disruption, and subsequent testing of the initially trained context (ABA) or of a different generalization context (ABC) should result in similar amnesia as observed to the generalization context used for reactivation (ABB) (Duvarci & Nader, 2004). In contrast with this prediction, we found that reactivation in generalization context B, which is clearly capable of retrieving the memory of context A as indicated by similar levels of conditioned responding in both contexts, does not result in amnesia when the later test takes place in context A. That is, we find amnesia in ABB-MDZ, but not in ABA-MDZ animals (Fig. 1). Whereas the lack of amnesia in ABA-MDZ subjects does not seem to fit with the reconsolidation account, advocates of that account may claim that our results do not provide conclusive evidence against it. Arguably, ABA recovery from amnesia could be due to incomplete memory reactivation of the original fear memory upon exposure to context B. Below, we will argue why this is unlikely to explain our results.

It is widely acknowledged that memory reconsolidation is stimulus selective. That is, non-reactivated elements of the original training situation are not assumed to destabilize and reconsolidate at the time of retrieval (e.g., Debiec et al., 2006; Debiec, Diaz-Mataix, Bush, Doyère, & LeDoux, 2013; Díaz-Mataix, Debiec, LeDoux, & Doyere, 2011). Building on this notion, one might argue that the lack of generalized MDZ-induced amnesia in ABA can be accounted for by incomplete reactivation of the original contextual fear memory through exposure to generalization context B. In other words, it can be argued that memory reactivation in generalization context B allows MDZ to exclusively disrupt the shared features of A and B associated with the US, whereas the association of other elements of context A with the US are preserved and would drive the recovery of fear when memory is tested in context A, in line with prior observations of stimulus renewal after extinction (e.g., Boddez et al., 2012; Vervliet, Vansteenwegen, Baeyens, Hermans, & Eelen, 2005).

The idea of partial memory destabilization of the original fear memory is interesting, but earlier results of our lab with reinforced memory reactivation challenge such hypothesis (Exp 4-5; Alfei et al., 2020). Others had already demonstrated that also exposure to the US (rather than to the unreinforced CS) can render associative fear memories vulnerable to post-reactivation amnestic treatments (e.g., Richardson et al., 1982). Based on those findings, it has been suggested that the presentation of the US during memory reactivation will render all elements originally associated with that US sensitive to disruption and reconsolidation in both rodents and humans (e.g., Dȩbiec et al., 2010; Dȩbiec et al., 2006; Deng et al., 2020; Dunbar & Taylor, 2017; Guo et al., 2020; Huang, Zhu, Zhou, Liu, & Ma, 2017; Liu et al., 2014; Luo et al., 2015; Thompson & Lipp, 2017; Xue et al., 2017). Adopting this logic, we found that reinforced reactivation in generalization context B (accompanied by a prediction error about the time of US arrival) indeed rendered the memory sensitive to MDZ, again resulting in amnesia in an ABB procedure, but not in animals tested in an ABA procedure (Exp. 4-5; Alfei et al., 2020). Together, these experiments indicate that direct associative retrieval of the original fear memory in an ABA procedure still does not induce generalized amnesia.

Further support against the idea that incomplete destabilization drives recovery of fear in the ABA experiments was obtained using an ABC experimental design (Exp. 6-7; Alfei et al., 2020). A, B and C contexts shared the same common features, in addition to unique features that were not shared with either of the other contexts. The reconsolidation account of forgetting would predict that MDZ after memory reactivation in B should result in amnesia at test in context C, given that there is no basis for fear responding in this context: the unique features of C have never been associated with the US, and the association of the common features of A, B and C with the US should have been disrupted by MDZ administration after memory reactivation in context B. Contrary to such prediction, no differences in freezing were observed between ABC-MDZ and ABC-SAL groups, and the amnesia found in ABB-MDZ was absent in ABC-MDZ animals.

Taken together, our earlier findings and the results presented here do not seem compatible with the predictions of the reconsolidation account of forgetting and pose a challenge for this account because (I) we show that ABB amnesia does indeed depend on destabilization of a previously acquired memory for A and displays other hallmark features of a destabilization-based phenomenon, yet (II) the lack of amnesia in ABA and ABC procedures (Experiment 1, 6; Alfei et al.,2020) is inconsistent with the notion that ABB amnesia is due to MDZ blocking the restorage of that destabilized memory for A. In addition, elsewhere, we have shown that (III) MDZ-induced ABB amnesia is sufficiently potent to resist rapid reacquisition and spontaneous recovery manipulations (see Alfei et al., 2020, Exp. 3 and 5). Persistence of drug-induced amnesia over time and insensitivity to recovery manipulations have traditionally been considered as behavioral indicators of permanent disruption of the engram (Elsey et al., 2018). The lack of amnesia in an ABA procedure, despite persistent amnesia in ABB, casts doubt on that logic as well. Rather than reflecting an irreversible blockade of reconsolidation, our results suggest that such amnestic procedures induce a (reversible) deficit in retrieval.

One way to salvage the reconsolidation account in light of our data would be to assume that retrieval of CFC through exposure to a generalization context B triggers consolidation of a GS-specific fear memory trace rather than destabilization-and-reconsolidation of the fear memory for the original context A. Previous research indeed suggests that the presentation of new contextual information at the time of memory retrieval can induce memory destabilization (e.g., Jarome, Ferrara, Kwapis, & Helmstetter, 2015; Winters, Tucci, & DaCosta-Furtado, 2009) but also initiate consolidation of a novel memory trace (e.g., Hupbach et al., 2008). The ability of hippocampal networks to perform pattern completion given input that only partially recapitulates prior stimulus input has been taken as support for their role in memory retrieval (e.g., Nakazawa et al., 2002; for review, see Rolls, 2013). These findings led to the hypothesis that a pattern completion mechanism can drive fear memory reactivation in the face of a generalization situation supported by the shared features of the initial learning situation (here, acquisition context A) and the retrieval situation (here, reactivation context B). At the same time, the discrepancies between the retrieval situation and the initial experience can trigger a pattern separation process that promotes the formation of a new fear memory trace. Post-retrieval injection of MDZ could then arguably disrupt the consolidation of that new contextual fear memory trace created for context B (and thus, the later expression of fear for B at test), rather than the reconsolidation of the original contextual fear memory for A. As mentioned earlier, consolidation and reconsolidation are both assumed to be temporally graded (Miller & Matzel, 2000). Therefore, the consolidation view could account for the lack of amnestic effect when MDZ administration is delayed for 6 h after reactivation (Experiment 2) and for the absence of amnesia on a STM test after memory reactivation along with the presence of amnesia on a LTM test in ABB MDZ-treated animals (Experiment 3).

Appealing as the idea that impaired memory consolidation accounts for MDZ-induced ABB amnesia may seem, the results of Experiments 4 and 5 render this explanation unlikely. Systemic and intracranial administration of IFEN prior to training is known to impair fear memory consolidation in rats (e.g., Bauer et al., 2002; Rodrigues et al., 2001; Sun et al., 2016). Experiment 4 showed that in our ABB procedure, IFEN administration prior to memory reactivation did not produce a memory impairment at test (compare the IFEN-SAL and SAL-SAL groups), in contradiction to the notion that exposure to B triggers the de-novo consolidation of a fear memory for context B. At the same time, IFEN did effectively block the post-retrieval amnestic effects of MDZ (compare the IFEN-MDZ and SAL-MDZ groups). Clearly, the lack of amnesia in the IFEN-SAL and IFEN-MDZ groups together with the amnesia observed in the SAL-MDZ group points towards destabilization of the contextual fear memory for A as a key mechanism underlying MDZ-induced ABB amnesia. Critically, further refutation that a consolidation-based process mediates MDZ-induced ABB amnesia comes from Experiment 5, where we found that MDZ at the dose used here, when administered shortly after acquisition, does not affect memory performance on a LTM test, suggesting that this dose of MDZ is incapable of interfering with the de-novo consolidation of a contextual fear memory.

Although studied for several decades now, it remains unresolved whether experimental post-retrieval amnesia reflects a failure of memory restorage (i.e., disruption of reconsolidation) or a failure to retrieve otherwise intact memory (Hardt, Wang, & Nader, 2009). The latter view proposes that post-reactivation pharmacological manipulations might alter the memory’s subsequent retrievability, effectively rendering it inaccessible at the time of test (Riccio, Millin, & Bogart, 2006). It appears that our findings are more readily reconciled with a retrieval deficit rather than with a restorage deficit. Similarly, early studies of the reconsolidation era demonstrated that post-retrieval amnesia induced by protein synthesis inhibitors can be reversed through presentation of the original US (e.g., Eisenberg & Dudai, 2004) and the passage of time (e.g., Lattal & Abel, 2004) (for review, see Urcelay & Miller, 2016). They are also in line with recent studies that used neurobiological probes to rescue forgotten contextual fear memories. (e.g., Roy, Muralidhar, Smith, & Tonegawa, 2017; Ryan, Roy, Pignatelli, Arons, & Tonegawa, 2015; for an elaborate overview, see Josselyn & Tonegawa, 2020) In particular, Ryan et al. (2015) found that although post-reactivation administration of anisomycin, a protein synthesis inhibitor, eliminated enhanced dendritic spine density and synaptic strength (considered key neural features of a consolidated fear memory trace), fear memory expression could be recovered by optogenetic activation of engram cells that had been active at the time of encoding. Findings from the same lab revealed that after blockade of memory consolidation, it is still possible to observe (protein-synthesis independent) changes in neural memory networks (Kitamura et al., 2017). Taken together, these findings indicate that even if a memory trace is modified by consolidation or reconsolidation mechanisms, memories remains retrievable.

In parallel with the abovementioned findings indicating that memories can be recovered after post-training and post-reactivation amnestic interventions, a number of observations in the last decade have suggested that pharmacologically induced post-retrieval amnesia can be reversed by re-administering the amnestic drug prior to testing (Briggs & Olson, 2013; Flint, Noble, & Ulmen, 2013; Gisquet-Verrier et al., 2015; Nikitin, Kozyrev, & Solntseva, 2019; Nikitin, Solntseva, & Nikitin, 2019; Rossato et al., 2015; Sierra et al., 2013). These findings are accounted for by the memory integration view of post-retrieval amnesia which postulates that information present around the time of retrieval (e.g., contextual features or an amnestic drug state) can get integrated into the memory representation (Gisquet-verrier & Riccio, 2018), such that successful retrieval of the memory may come to depend on the presence of that information. As a result, the more the circumstances at the time of test resemble those of reactivation, the less amnesia should be observed (Tulving & Thomson, 1973). Clearly, the drug state induced by MDZ can be one example of such a state; physical aspects of the experimental context are another obvious example.

If that logic is applied to our current and earlier findings, it seems reasonable to assume that, other things being equal, after reactivation using context B, memory should be more readily expressed in ABB than in ABA and ABC, given that a test in context B (even in the absence of MDZ) recreates the circumstances at the time of memory reactivation better than a test in C (ABC) or A (ABA). Yet, as discussed in Alfei et al. (2020, Exp. 3), we obtained weaker memory expression in ABB-MDZ than in ABA-MDZ, as also shown in Exp. 1 of the current paper. Likewise, memory expression should be expected to be at least as strong in ABB as in ABC according to the memory integration account, in contrast to what we observed in Alfei et al. (2020, Exp. 7). Furthermore, the memory integration view predicts more amnesia in AAB-MDZ than in ABB-MDZ, whereas we observed similar levels of amnesia in both (Alfei et al., 2020, Exp. 3). In conclusion, central findings of our current and earlier work, while generally consistent with a retrieval deficit view, are problematic for the memory integration view of post-retrieval amnesia.

Finally, it is worth noting that, although some of the earliest evidence supporting state-dependent memory retrieval comes from studies using GABA-A agonists, e.g., amobarbital, alcohol or diazepam (e.g., Goodwin, Powell, Bremer, Hoine, & Stern, 1969; Ley et al., 1972, but see Meyer et al., 2017) others have shown that disruption of Pavlovian fear memory by MDZ is not due to state dependency (e.g., Harris & Westbrook, 1998), impaired locomotion, anxiolytic properties or sedative effects (e.g., Pain et al., 2002). Therefore, the assumption that the observed failure to retrieve the memory is due to the lack of MDZ prior to test seems highly disputable

Even if the memory integration account of post-retrieval amnesia cannot readily explain our findings, it would remain possible that a different form of retrieval interference is at play. In particular, one might argue that the memory reactivation session in generalization context B functions as an extinction session, the effect of which could be facilitated by the anxiolytic effect of the benzodiazepine MDZ (Hart, Harris, & Westbrook, 2009, 2010). Extinction is considered to be a new active learning process, leading to formation of a CS-no US inhibitory trace that competes for retrieval with and temporarily suppresses the original CS-US association (Bouton, 2004). It could be argued that MDZ’s anxiolytic effects may facilitate the acquisition of this specific inhibitory association to the reactivation context, resulting in a reduction of fear when animals are later tested in that context again (AAA or ABB). Yet, there are a number of arguments that go against such interpretation. First, studies of Hart, Harris and Westbrook (2009, 2010) – all conducted in a single context – only observed a facilitating effect of MDZ when it was administered prior to the start of a second extinction session which had been *preceded* by an initial extinction session one day earlier. In contrast, inhibition of contextual fear was impaired when MDZ was injected prior to the first extinction trial. The authors argued that the drug’s anxiolytic effects during the second extinction session reactivated and strengthened the previously acquired inhibitory memory. It seems unlikely that there would be such a facilitation in our experimental design, where MDZ is administered after a single non-reinforced reactivation session of 2 min. Additionally, it is worth noting that we have previously reported that a brief reinforced reactivation session in generalization context B (in which the timing of the US was altered relative to acquisition) followed by MDZ also resulted in attenuated fear memory expression (see Alfei et al., 2020, Experiment 4). It is unclear how such reinforced reactivation could engage the formation of an inhibitory memory trace that would be facilitated by MDZ. Second, if the amnesia we observed were indeed due to MDZ-facilitated inhibitory learning, one should expect such amnesia to be lifted by experimental manipulations such as reinstatement or spontaneous recovery (Bouton, 2004). In contrast, we have found attenuated reinstatement and spontaneous recovery in ABB MDZ-treated rats (see Alfei et al., 2020, Experiments 2-5). Third, rather than facilitating the formation of an inhibitory memory trace, it has been reported that post-extinction administration of benzodiazepines counteracts extinction learning, which makes sense given the fact that inhibitory memory formation depends on reduced GABAergic transmission after extinction training (e.g., Bustos et al., 2006, for reviews, see Makkar, Zhang, & Cranney, 2010; Myers & Davis, 2002). Likewise, we previously found (in an AAA design) that after an extended non-reinforced memory reactivation of 15 min (i.e., extinction), systemic MDZ blocked the formation of inhibitory learning, with MDZ-treated rats showing more freezing than controls at test, again in contrast with a facilitation of inhibitory learning (Ferrer Monti et al., 2017, Experiment 2; also see Alfei et al., 2015, Experiments 7A and 8A). Fourth, if MDZ caused the animals to acquire a context-specific inhibitory association, this cannot readily explain why we observed a long-lasting memory deficit in AAB MDZ-treated animals (see Exp. 3, Alfei et al., 2020), as the test took place in a different context than the one where the inhibition was generated (and presumably facilitated). In other words, the results in AAB should then be similar to what is found in ABA and ABC experiments, which is not the case. Taken together, the above-mentioned arguments suggest that facilitation of an inhibitory memory by MDZ cannot easily account for the post-retrieval diminished fear expression induced by MDZ that we observed.

Regardless of which specific account of amnesia can best explain the whole of our results (if any), the observation that post-retrieval amnesia can be reversed is more readily reconciled with the general idea of amnesia reflecting a retrieval deficit than with the notion of a disruption in memory restorage. One outstanding issue for such a retrieval deficit view of post-retrieval amnesia, however, concerns the conditions under which a reactivated memory enters into a transient state of sensitivity to retrieval interference. According to the retrieval deficit view of amnesia, memory reactivation can place the memory in a malleable state. Yet, advocates of such view of post-retrieval amnesia might agree with the statement that memory reactivation *per se* is not sufficient to render the memory sensitive to retrieval interference. Likewise, advocates of a storage deficit view of amnesia support the notion that only under certain conditions a reactivated memory can be destabilized and reconsolidated (e.g. Díaz-Mataix et al., 2013; Exton-McGuinness, Patton, Sacco, & Lee, 2014; Pedreira et al., 2004). Our data clearly indicate that post-reactivation amnesia is obtained only when specific conditions are fulfilled at the time of memory reactivation (also Alfei et al., 2015, 2020). Historically, retrieval deficit accounts of post-reactivation amnesia have paid little attention to those conditions, which have classically been linked to the initiation of a memory destabilization process.

The absence of ABA and ABC amnesia as opposed to ABB amnesia (as observed in Experiment 1 and elsewhere (Alfei et al., 2020)) clearly does not support a restorage deficit view of post-reactivation amnesia, that is, it does not fit with the notion that a reactivated memory trace can be rewritten by destabilization-and-reconsolidation processes and rather indicates that post-reactivation amnesia results from a drug-induced retrieval deficit. Still, our findings (Experiments 1-4; see also Alfei et al., 2020, Experiment 4) indicate that for such retrieval deficit to occur following reactivation, conditions need to be met that give rise to memory sensitivity to interference. A retrieval deficit view of amnesia needs to accommodate the data presented here and allow for the observation that reactivation-induced sensitivity to interference requires NMDA receptor activity (see Experiment 4) and an optimal degree of prediction error during memory reactivation (Alfei et al., 2020, Experiment 4), is transient (Experiment 2), and is not expressed on an immediate test after reactivation (Experiment 3). Accommodating those observations will likely require the incorporation of some elements of classic reconsolidation theory into a retrieval deficit account, such as the notion of engram destabilization (Pascale Gisquet-Verrier & Riccio, 2012).

To recap, we here investigated the characteristics of MDZ-induced ABB post-retrieval amnesia using different behavioral and pharmacological approaches firmly established in the reconsolidation literature. Our data indicate that such ABB amnesia cannot be accounted for by impaired consolidation of a separate fear memory for B. ABB amnesia does display the classic features of a destabilization-and-reconsolidation mediated phenomenon that have been established for regular AAA amnesia (Finnie & Nader, 2012; Nader & Hardt, 2009). However, the reversible nature of this ABB MDZ-induced amnesia defies a reconsolidation blockade explanation of post-reactivation amnesia. Whether our findings with systemic administration of MDZ can be generalized to other amnestic agents such as protein synthesis inhibitors (e.g., anisomycin, rapamycin; for reviews, see Jarome & Helmstetter, 2014; Tronson & Taylor, 2007), warrants further investigation. Critically, however, if translation from basic science to interventions is considered an important goal of drug-induced amnesia research (Elsey et al., 2018; Elsey & Kindt, 2017; Faliagkas, Rao-ruiz, & Kindt, 2018; Kindt, 2018), there is an important caveat to be deduced from our results. The observation that techniques of pharmacological amnesia induction durably prevent the re-emergence of fear memory expression has been taken to indicate the superiority of reconsolidation interference over other techniques (e.g., exposure treatment) as a means of therapeutic forgetting (Beckers & Kindt, 2017). The present findings give cause to question the utility of reconsolidation-based treatments, given the failure to broadly attenuate generalized threat responding after memory reactivation using a generalization context (see the lack of amnesia in ABA and ABC). A crucial challenge for future basic and (pre)clinical research indeed pertains to the development of therapeutic approaches that can more successfully tackle maladaptive fear memory generalization.

## Author Contributions

Conceptualization: J.M.A., T.B., L.L., D.D.B. Methodology and design: J.M.A., T.B., L.L., Validation: J.M.A., T.B., L.L. Data analysis: J.M.A., H.D.G. Investigation: J.M.A., H.D.G. Dara curation: J.M.A., H.D.G. Writing – Original Draft, J.M.A., T.B., L.L. Writing – Review & Editing: J.M.A., T.B., L.L., D.D.B. Visualization: J.M.A. Supervision: T.B., L.L. Project administration: T.B., L.L. Funding acquisition, T.B.

## Funding sources

This work was supported by a Consolidator Grant of the European Research Council (ERC) [T. Beckers, grant number 648176]

## Declarations of interest

The authors declare no competing interests.

## Notes

https://osf.io/zyx3p/

### Competing Interest Statement

The authors have declared no competing interest.

